# Organization, Neuronal Composition, and Dynamics of Neuronal Ensembles in the Songbird Auditory Forebrain

**DOI:** 10.64898/2026.01.19.700427

**Authors:** Felipe A. Cini, Luke Remage-Healey

## Abstract

Auditory learning is a key component of vocal learning and communication. Neurons involved in auditory learning are typically examined as single encoders, but there is increasing evidence that the coincident activity of groups of neurons, or ‘ensembles’, is important for the processing and transmission of sensory information. In songbirds, a forebrain region analogous to mammalian secondary auditory cortex, the caudomedial nidopallium (NCM), is crucial for representing and learning auditory signals. In the awake state, NCM neurons exhibit stimulus-specific adaptation in response to repeated presentation of song stimuli, considered a form of auditory working memory. Yet, how complex stimuli like song are encoded by networks of excitatory and inhibitory neurons is essentially unknown. Using *in-vivo* single-unit electrophysiology, we systematically unveiled neuronal ensembles that operate at high temporal precision (< 10 ms) across the NCM of awake zebra finches (*Taeniopygia guttata*). Our data show that multiple ensembles are concurrently activated by song with high temporal precision, and that their neuronal composition is heterogeneous, topographically proximate, and biased towards excitatory members. NCM ensembles adapt to song playback and, notably, become more stimulus selective over tens of minutes, accompanied by fast remodeling (membership gain and/or loss) during adaptation. Altogether, our results suggest that song representations in the forebrain can be conveyed by multiple, dynamic network ensembles in parallel. These findings advance our knowledge of the composition, dynamics, and neuronal network reorganization of ensembles as complex sensory stimuli become increasingly familiar.

## Introduction

Much of our understanding of cognitive processes, such as sensory processing, memory, and information integration arises from studies of single neurons. Yet, the recurring patterns of coordinated activity of groups of neurons – neuronal ensembles – have long been proposed to play a crucial role in these processes ^1,2^. New technologies have allowed for the detection of a large number of neurons in a single recording, enabling the identification and systematic study of neuronal ensembles. Local ensembles, detected through correlated neuronal activity, have been shown to effectively encode information beyond the capability of single neurons ^3–8^. Additionally, recent in-silico modeling suggests that correlated neuronal activity allows the scaling of memory capacity and nonlinear computations ^9^. Understanding these emergent properties may provide critical insights into how networks of neurons encode and store information. However, key ensemble properties remain largely unknown, including their cellular composition, stability, selectivity, and optimal synchronization time between members.

Inhibitory interneurons play a key role in generating low (20-30Hz) and high (30-100Hz) gamma oscillations and promoting local network synchronization, while at the same time constraining noise correlations (Cardin et al., 2009; Sohal et al., 2009; Veit et al., 2017; Chen et al., 2017). Most studies of network ensembles focus exclusively on principal/excitatory cells and/or do not differentiate ensemble membership by cell type ^10,11^. Therefore, there is a significant gap in understanding the contributions of inhibitory vs excitatory cell types to ensemble membership and their interactions.

Studies of neuronal ensembles have been mostly gleaned from rodent species, but we now have an opportunity to expand this specific reference frame. Songbirds, especially zebra finches (*Taeniopygia guttata*), offer valuable insights into auditory processing and learning. Zebra finches rely heavily on vocal communication for various purposes, including pair bonding, individual recognition, and mating ^12,13^. The caudomedial nidopallium (NCM), a region of the secondary auditory pallium in songbirds, plays a key role in auditory stimulus categorization, learning, and novelty detection (Chew et al., 1995; Jarvis et al., 1995; Lu & Vicario, 2017; Macedo-Lima & Remage-Healey, 2020; Ribeiro et al., 1998; Schneider & Woolley, 2013). Studies in NCM largely focus on single-unit activity encoding of sounds or sound features ^14,19–22^, and excitatory and inhibitory neurons in NCM can be distinguished unambiguously by their electrophysiological waveforms in vivo and ex vivo (Spool et al., 2021). Inhibitory neurons show stronger sound-evoked responses, higher firing rates, and have lower stimulus selectivity than excitatory neurons ^23–27^, matching similarly divergent response properties in mammals. Thus, understanding ensemble activity in this system can help uncover broader principles of auditory processing and communication in a vocal learning species.

A characteristic feature of NCM neurons is their fast stimulus-specific adaptation (SSA) when presented repeatedly with the same auditory stimulus ^14–16^. This adaptation represents a form of cellular memory in which neural responses adjust dynamically to reflect recent stimulus-related information and experience. In rodent cortex, circuit dynamics during SSA show suppression of phase coupling between spikes and local field potential in the beta range ^28^, and neuronal ensembles that detect novel, non-adapted stimuli ^29^. Furthermore, two distinct classes of inhibitory neurons, parvalbumin (PV) and somatostatin (SST), play different roles in SSA, with SST facilitating adaptation ^30,31^. However, the dynamics of ensemble behavior during SSA remain unclear in any system. We hypothesize that neuronal ensembles undergo rapid remodeling during adaptation, retaining efficient encoding for familiar stimuli, and updating their representation to enable rapid stimulus recognition in a cell-type specific manner.

To address how neuronal ensembles contribute to auditory information encoding, we examined their stability during SSA in NCM. Recent studies have described groups of neurons as ‘ensembles’ collected statistically across recordings separated in time from multiple animals, providing glimpses of the role of different cell types to songbird population auditory processing and decoding ^32,33^. Here, we build on these advances by using high-density silicon recording probes to simultaneously record dozens of neurons and isolate concurrently-activated neuronal ensembles. In this way, we can now investigate network ensemble adaptation and stability to gain insights into the mechanisms underlying auditory plasticity and memory. Our study also explores principles of ensemble composition - particularly the role of inhibitory vs. excitatory neurons - and the temporal precision of neuronal synchronization. These findings show that ensembles undergo fast remodeling during SSA, that ensemble membership is biased toward excitatory units, and that they synchronize within an ultra-short, six-millisecond timescale.

## Methods

### Animals

Male and female adult zebra finches (n = 8, 4 females, >90 days old) from the University of Massachusetts Amherst colony were used for the experiment. Birds were not actively breeding during the experiments (single-sex cages) and were on a 14:10 hour light-dark cycle. All procedures were in accordance with the Institutional Animal Care and Use Committee at the University of Massachusetts Amherst (Protocol #2019-0058).

### Surgery and Electrophysiological recording

Birds were food-deprived for one hour before surgery, anesthetized with isoflurane (1-2% in O_2_), and then placed on the stereotaxic apparatus (Kopf) with the head tilted at a 45° angle. Bilateral craniotomy was performed at markings 0.9mm anterior and 0.7mm lateral respective to the bifurcation of the midsagittal sinus, and then the meninges were resected. A small craniotomy on the anterolateral part of the skull for the implantation of a silver ground wire using cyanoacrylate and dental cement. Craniotomies were sealed with Kwik-Cast (World Precision Instruments). A custom-made headpost was lowered on the beak and fixed to the skull with dental cement. Recordings from both hemispheres were performed 1-3 days after surgery.

For the electrophysiological recordings we used open source, 64-channel silicon probes (model 64G) from the Masmanidis Lab – UCLA ^34^. Electrode impedance was set between 300-500 kOmhs and confirmed on the day of the recording before insertion. Birds were awake restrained and head-fixed for the entirety of the recording. After the bird was secured, the Kwik-Cast was removed and the probe was inserted medio-caudally in register with markings. To confirm probe location, the probe was briefly dipped in DiI (Diluted in 100% ethanol) and allowed to dry before descent ∼1.6mm from the brain surface.

The probes had 64 channels on two shanks, 300µm apart, with 32 channels each. The channels within one probe were arranged along a 525µm long axis, and the space between two contact points was 25µm vertical spacing and 20µm horizontal spacing. Recordings were amplified and digitized by a 64-channel amplifier and evaluation board (RHD2000 series; Intan Technologies) and sampled at 30 kHz using Open Ephys software. An Arduino Uno (Arduino) was connected to the recording computer to deliver TTL pulses to the evaluation board’s DAC channel bracketing the beginning and end of the audio stimuli (described below) to optimize detection during analysis. Audio playback and TTL pulses were controlled by a custom-made MATLAB (MathWorks) script, which also controlled the Arduino and sent a copy of the audio analog signal to the evaluation board ADC channel. Each bird underwent one recording per hemisphere per day. Following recordings, birds were transcardially perfused with paraformaldehyde 4% in 0.025 M phosphate. Brains were extracted and placed in 30% sucrose and stored in -80 ^°^C.

### Stimuli

32 song files from unrelated birds were randomly and equally split into four sets, containing 8 songs each. For each animal, one set was used per hemisphere (search stimuli to locate NCM were non-songs). This ensured that each bird heard novel stimuli to avoid long-term adaptation in NCM firing states across sessions and days (Chew et al., 1995; Ribeiro et al., 1998; Vates et al., 1996). Therefore, a novel stimulus set was used for each recording to control for familiarity effects.

Each stimulus set consisted of five conspecific songs, two heterospecific songs, and a white noise pulse of typical song duration, repeated 25 times each in pseudorandomized order. The interstimulus interval was pseudorandom within the interval 5 +- 2s. The speaker was positioned 30 cm in front of the animal, equidistant to each ear. The sound level was amplified to peak ∼65 dB as measured by a sound level meter at the animal’s position (RadioShack). Playback trial duration lasted around 25 min. Recordings were performed inside a Faraday cage inside an anechoic booth (Industrial Acoustics). Recordings were made from each hemisphere on the same day or 1 day apart. For each animal’s first recording, the starting hemisphere was randomized and then counterbalanced between sexes. The stimulus set was also initially randomized, then counterbalanced across sexes and hemispheres, but the subset selected for each was always randomized.

### Electrophysiology Analysis

Single-unit sorting was done with Kilosort4 ^36^. Sorting results were manually curated in Phy (https://github.com/cortex-lab/phy), and well-isolated units were used (high signal-to-noise ratio; low violation of refractory period, low contamination with other units; segregation in waveform PCA space). After sorting (mean = 54.35 units detected per recording site, SD = 14.9), 2000 waveforms were selected randomly for each single unit. The waveform features were aligned, then averaged, and their waveform features were measured (peak-to-peak duration and ratio) in MATLAB.

Normalized z-score evoked responses to each stimulus were calculated by the formula:

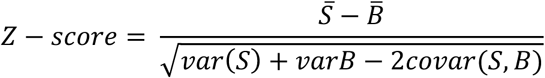

Where S and B are the stimuli and spontaneous firing rates across stimulus trials, respectively. After computing the z-score by stimulus, those were averaged to yield a single z-score per unit per stimulus type (conspecifics, heterospecific, white noise). Adaptation rates were calculated using trials 6–25, which is the most linear phase of the adaptation profile in NCM ^37,38^. For each stimulus, the stimulus firing rate across trials was normalized by the firing rate on trial 6. Then, a linear regression was calculated between trials 6 and 25. For each stimulus type, the average adaptation slope across stimuli was used for each unit.

### Ensemble detection

We identified neuronal ensembles by combining principal component analysis (PCA) and independent component analysis (ICA) ^39^. Spike times for the whole recording were binned, creating a spike matrix where each row represented the activity of a single-unit and each column indicated the number of spikes within each time bin (Supplementary figure 1a). The activity of each unit was then normalized through z-scoring, followed by calculating the correlation matrix over the normalized spike matrix for all pairs of neurons.

To determine the number of ensembles in each recording, we applied a PCA to the z-scored spike matrix to obtain the eigenvalues. Eigenvalues that were higher than the upper limit of the Marcenko-Pastur distribution ^40^ were considered to be the result of significantly correlated activity. We then applied a fast ICA over the eigenvectors of the significant eigenvalues. The resulting independent components (ICs) of the fast ICA represent each neuron’s contribution to its respective ensemble. Ensemble activation strength was calculated by projecting the Ics back onto the z-scored spike matrix ^39^.

Neurons whose IC weight exceeded the mean weight of the ensemble by ±1.5 standard deviations from the distribution of all IC weights were considered members of the ensemble. The threshold for considering an ensemble to be activated was 3 standard deviations of the activation strength distribution.

Evoked activation z-scores and adaptation profiles for the ensembles were created using a similar calculation as above, with single neurons z-score evoked firing rate and adaptation; however, instead of using spikes the activation of the ensemble was defined by the 99.9^th^ percentile of the activation strength distribution. A limitation of this approach is that the PCA can only identify linearly separable components and the ICA assumes that the components are non gaussian. Even with these liminations, the ensemble identification strategy we employed is most commonly adopted and has been used in a variety of systems and species ^41,42,42–45^.

### Measuring ensemble remodeling/stability

To identify the extent to which ensembles undergo remodeling during SSA, we measured the trend of activation strength of the ensemble throughout the recording using the non-parametric Mann-Kendall trend test ^46^. If the p value was below 0.05 and the trend was negative, we classified the ensemble as “remodeling”, if the p value was below 0.05 and the trend was positive it was classified as “strengthen connectivity”, and if the p value was higher than 0.05, it was classified as a “stable ensemble”.

One concern with the above analysis scheme is that it only acquires ensemble membership once integrated across the entire recording, perhaps leading to an over-estimation of ensemble stability. To address this, we segmented the recording in half and performed ensemble detection on the two halves separately. After detecting the members of the ensembles, we compared their composition in the first and second half using the Jaccard Similarity Score (Pérez-Ortega et al., 2021). The scores range from 0, no overlap between members, to 1, total overlap of members.

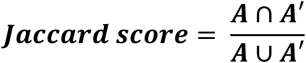

- A: Members of an ensemble in the first half of recording.
- A’: Members of an ensemble of the second half of the recording.

We ranked all compared pairs (first and second half ensembles) based on the Jaccard score, the pair with the highest score was then used for evaluation. If the score was below 0.3 (≅ 50% similarity), then they would be considered different ensembles (the ensemble detected in the second half was considered a completely new ensemble) ^47^. If the score was above 0.3 but below 0.9, the ensemble was defined as remodeled, meaning that they are still the same ensemble but have lost, gained, or changed members. Lastly, if the ensemble pairs had a score above 0.9, it was considered a stable ensemble.

### Bayesian decoder

To help identify the best time bin for ensemble detection of individual sorted units, we also used Bayesian decoding ^48–51^:

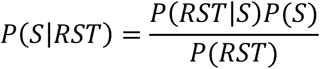

Where:

- S: stimulus
- RST: response strength for each stimulus playback calculated as the difference in spikes between pre- and during playback.
- P(S|RST): Posterior probability of stimulus S given the ISI.
- P(RST|S): Likelihood of observing the ISI given stimulus S.
- P(S): Prior probability of stimulus S
- P(RST): Evidence of RST

### Mutual information

To help disambiguate between optimal time bin for ensemble synchronization, mutual information (MI) was calculated between the evoked response RST and stimulus for each ensemble and bin size. MI was computed using the *mutual_info_regression* function from the scikit-learn package in python ^52^. This function relies calculates MI based on the entropy estimation from k-nearest-neighbors distance ^53^. This approach does not assume linearity or normal distribution. The resulting MI values reflect the amount of stimulus-related information carried by each ensemble for each temporal bin-size.

### Statistical analysis

Statistical analyses were performed in Python ^54–57^. To compare unpaired groups, we used general linear mixed models. For single unit analysis, the GLM model included stimulus type, cell type, sex and hemisphere, while bird ID was considered a random effect. For ensemble level analysis, the GLM model included stimulus type, and sex, while hemisphere and bird ID were considered random effects. For post hoc analysis we used the Tukey test. To determine if two samples are drawn from the same distribution, we used the Kolmogorov-Smirnov (KS) test.

## Results

### Single unit activity in accordance with prior reports

We performed electrophysiology recordings from the caudomedial nidopallium (NCM), the equivalent of the secondary auditory cortex in mammals, using 64-channel silicon probes during the playback of conspecific songs, heterospecific songs (Bengalese finch – *Lonchura striata domestica*), or white noise (Figure 1a). Well-isolated single units were classified as either inhibitory (ptp duration <0.35 ms, Figure 21a) or excitatory (ptp duration ≥ 0.35 ms) based on the average waveform (2000 samples) of that unit (Figure 1b). We note that Spool and colleagues (2021) determined the molecular identity of extracellular waveforms in NCM to excitatory vs. inhibitory neurons using optotagging and a similar ptp duration threshold. Out of all units recorded (Birds = 8, Recordings = 14, single-units = 759), the majority were excitatory (excitatory: mean = 67.23%, sd = 13.08%, inhibitory: mean = 32.77%, sd = 13.08%; Figure 1a). Cells in the NCM are auditory responsive, and there was an interaction between response by cell type and stimulus type (ý= -0.204, se = 0.38, z = -5.33, p < 0.001; Figure 1e). Inhibitory units responded more strongly to conspecific than heterospecific songs (*mean difference* = -0.41, 95%CI [-0.74, -0.07], p =0.006), while excitatory units responded equally strongly to both song types (*mean difference* = -0.04, 95%CI [-0.25, -0.16], p =0.987). Furthermore, inhibitory units showed an overall greater evoked response to auditory playback than excitatory units (ý= -0.613, se = 0.027, z = -22.98, p < 0.001). Next, we examined the units’ selectivity based on their response percentage to different conspecific songs, finding that excitatory units were more selective than inhibitory units (ý= -11.86, se = 0.92, z = -13.67, p < 0.001). These findings are in general accordance with prior electrophysiological studies of excitatory vs inhibitory neurons in NCM ^24,25,58–61^.

**Figure 1.**
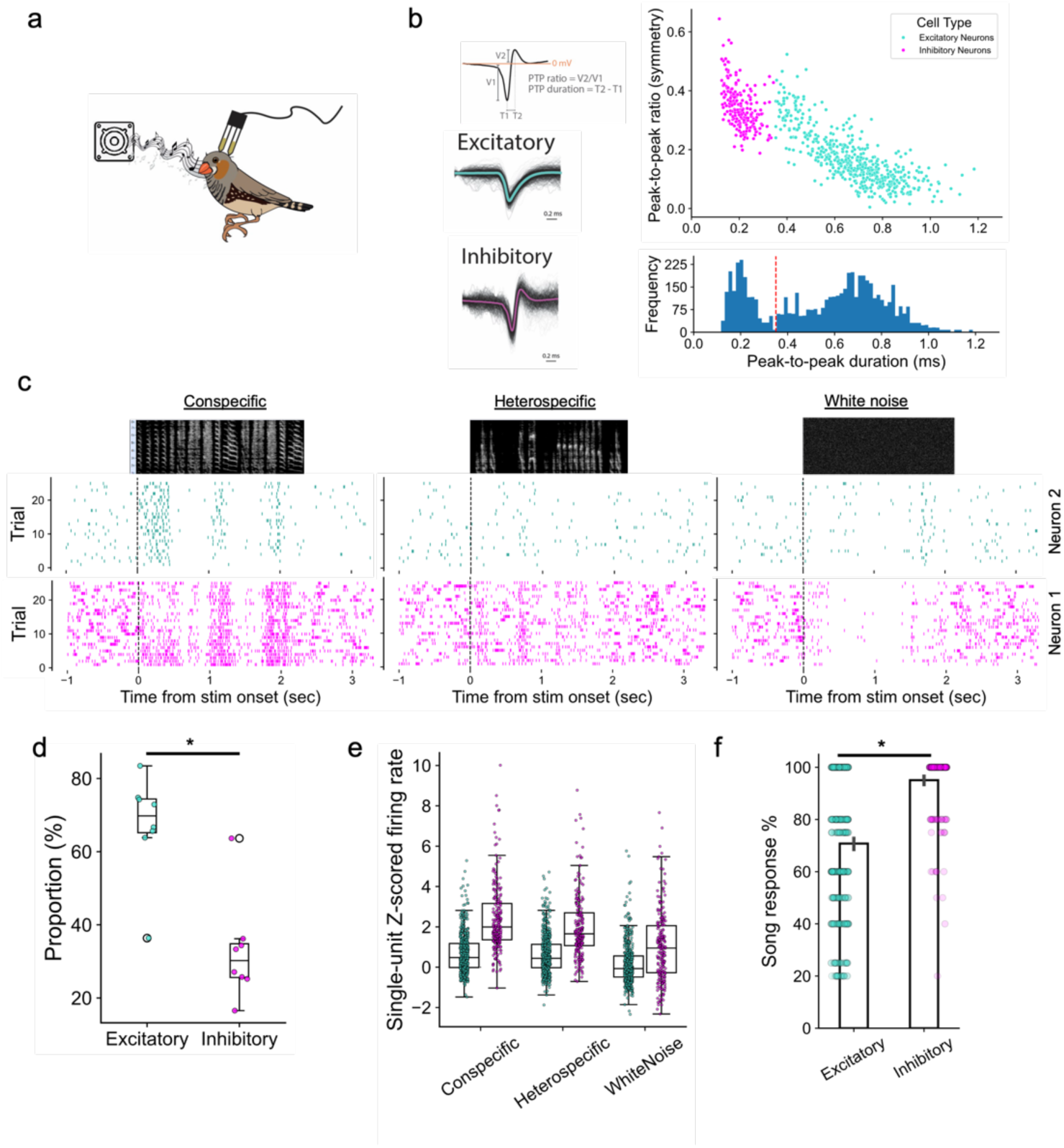
Excitatory and inhibitory neurons respond differently to songs. a) Experimental scheme for the electrophysiology recording in head-fixed awake zebra finches. **b)** Representative spike traces for one inhibitory and one excitatory cell. Scatterplot with the peak-to-peak (ptp) duration and ratio for all detected neurons (n = 15 recordings, from 8 birds - 4 males, 4 females - and 759 neurons), excitatory and inhibitory neurons were differentiated by the ptp duration, cells were defined as inhibitory if ptp < 0.35 ms. **c)** Representative PSTHs from one inhibitory and one excitatory cell in response to different stimuli. **d)** Proportion of excitatory and inhibitory cells per recording. **e)** Evoked normalized response to the different stimulus types. Tukey’s post hoc test results are shown at the bottom. * Significant values p <0.05. **f)** Selectivity measured as percentage of songs each unit responded to; 100% = all songs, 0% = none.

### Ensemble detection and composition

After verifying reliable single-unit activity, we investigated how these units work together to process auditory information by examining coordinated neuronal ensemble activity (Figure 2a). Here, we define an ensemble as a group of neurons recorded simultaneously from the same recording site that fire together within a short (< tens of ms) timeframe. To detect ensembles (Figure 2b, c), we binned the data in a selected time bin (see below) and then normalized the firing rate of each unit. We then calculated a correlation matrix between all units and performed a principal and independent component analysis on this matrix (Supplementary Figure 1).

**Figure 2.**
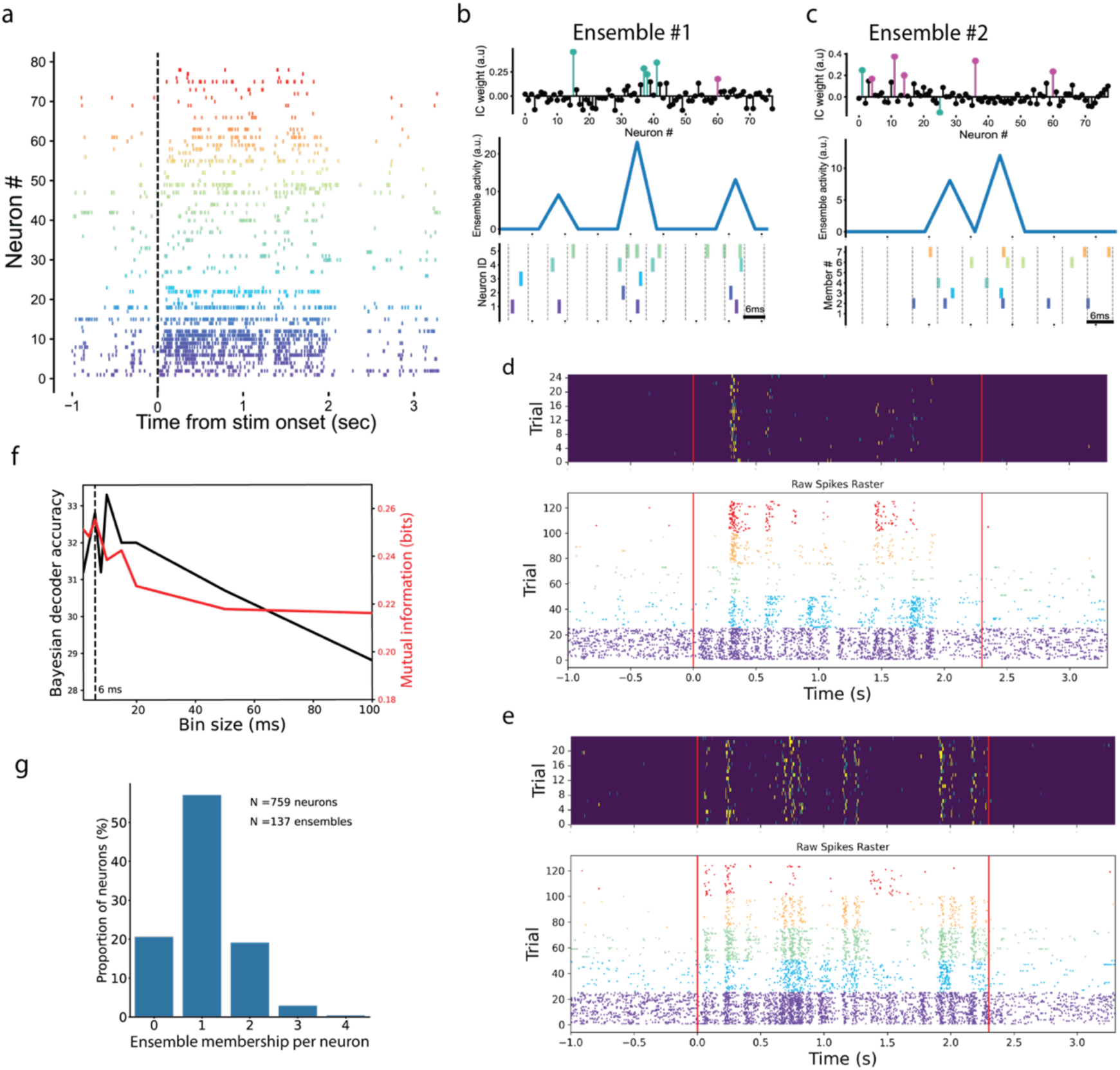
**Neuronal ensemble detection and response**. **a)** Example of neuronal response to stimuli from one recording. Each color represents a different neuron. **b**,**c,)** Examples of detected ensembles. *Top:* shows the contribution each neuron cell ensemble IC weights to ensemble membership (black circles: non-members, pink: inhibitory ensemble members, light blue: excitatory ensemble members. *Middle:* Shows ensemble activation strength followed by *(Bottom)* spiking of individual ensemble members. **d, e)** *Top*: heatmap of the ensemble 1 activation for 25 trials for two different songs. *Bottom:* raster plot for the 5 different neurons that make up the ensemble. **f)** Optimal bin size of 6 ms selected based on maximum mutual information and decoder accuracy. **g)** Proportion of neurons that are members of 0, 1, 2, 3 or 4 ensembles.

In the current literature, the synchronization timeframe for neurons in an ensemble has not been fully settled. Therefore, we tested a series of bin sizes to identify the optimal duration for ensemble detection and function. Although not statistically significant, the 6ms bin size had the highest mutual information (MI) (Figure 2f) and the second highest accuracy with the Bayesian decoder (Figure 2f). Consequently, we performed further analyses using the ensembles detected with the 6ms bin size. Of the 759 neurons detected, most participated in at least one ensemble; 57.05% participated in only one ensemble,22.40% participated in multiple ensembles, and 20.55% were not part of any ensemble (Figure 2g). The identified ensembles with this time bin were also auditory responsive and exhibited pronounced temporal precision (Figure 3a, b), showing greater evoked response to songs than to white noise (β= -1.093, se = 0.112, z = -9.75, p < 0.001; Figure 3c). NCM ensembles were as song selective as excitatory units but more selective than inhibitory units (Figure 3d). Ensembles also showed distinct, non-overlapping response patterns to individual song stimuli (Figure 2d,e; Figure 3 a,b) with strikingly narrow (< 20 ms) activation epochs. Thus, in principle, ensembles are an emergent feature among member neurons for highly-refined, song-selective response properties beyond that observed in the activity of single NCM neurons.

**Figure 3.**
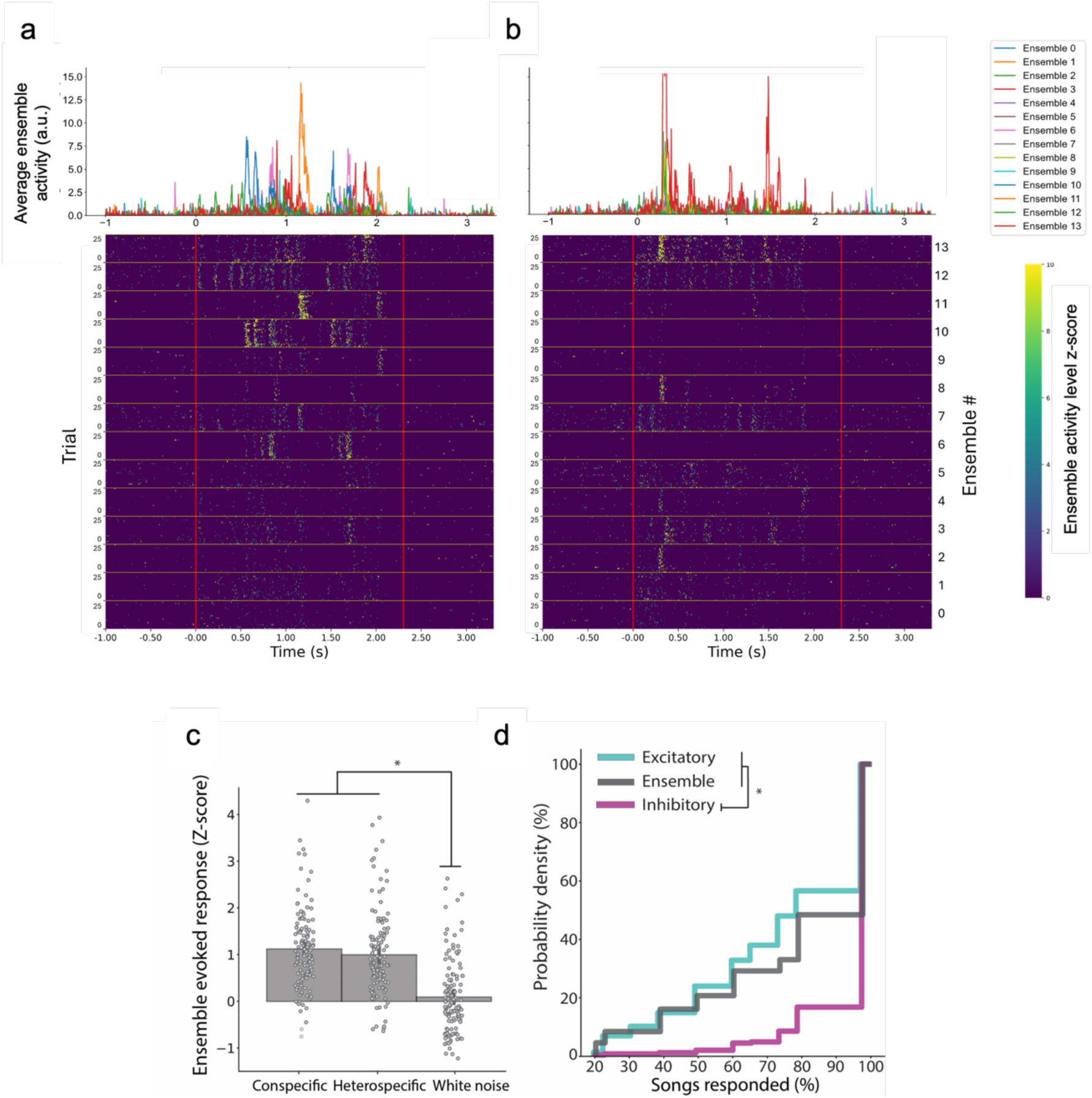
Ensembles respond selectively to songs. Ensembles respond differently to different songs, with ensembles showing distinct response patterns to the same stimulus. **a, b)** *Top*: overlapped average activation strength for different ensembles in response to songs. Different ensembles are displayed with different colors. *Bottom:* heatmap for the 25 trials for different ensembles separated by the yellow lines. **c)** Evoked normalized response to the different stimulus type. Tukey’s post hoc test. **d)** Selectivity measured as percentage of songs each unit/ensemble responded to; 100% = all songs, 0% = none. Ensembles and excitatory cells are more selective than inhibitory. * p-value <0.05.

What is the topographic and cellular composition of ensembles? We characterized the composition and distribution of the members of detected NCM ensembles and compared them to non-comembers (defined as neurons that are not members of the same ensemble) and to members of randomly shuffled ensembles, using electrode location as a proxy for unit location. Ensemble members were physically closer to each other than randomly defined ensembles (Figure 4c). The maximum distance between two members was also shorter than that between non-members and also than those in the random distribution, particularly within < 100μm recording separation distance (Figure 4d). We also analyzed ensemble composition, finding that most were heterogeneous and biased toward excitatory units when compared to randomly generated ensembles (ks = 0.21, p < 0.001; Figure 4b). Although the recorded NCM neurons were on average 32.77% inhibitory and 67.23% excitatory, exclusively excitatory ensembles comprised 30% of the population, while 4.75% were exclusively inhibitory (Figure 4b), and most were mixtures of both inhibitory and excitatory. Also, ensemble composition scaled with time bin; we found that the proportion of excitatory members increased with longer time bins (Suppl. Figure 2). In summary, we observe that ensembles in the songbird auditory pallium are drawn from proximate (<100 µm) neighbors, that ensembles synchronize optimally in an ultra-short time frame of 6ms, and that ensemble membership is prominently biased towards excitatory members.

**Figure 4.**
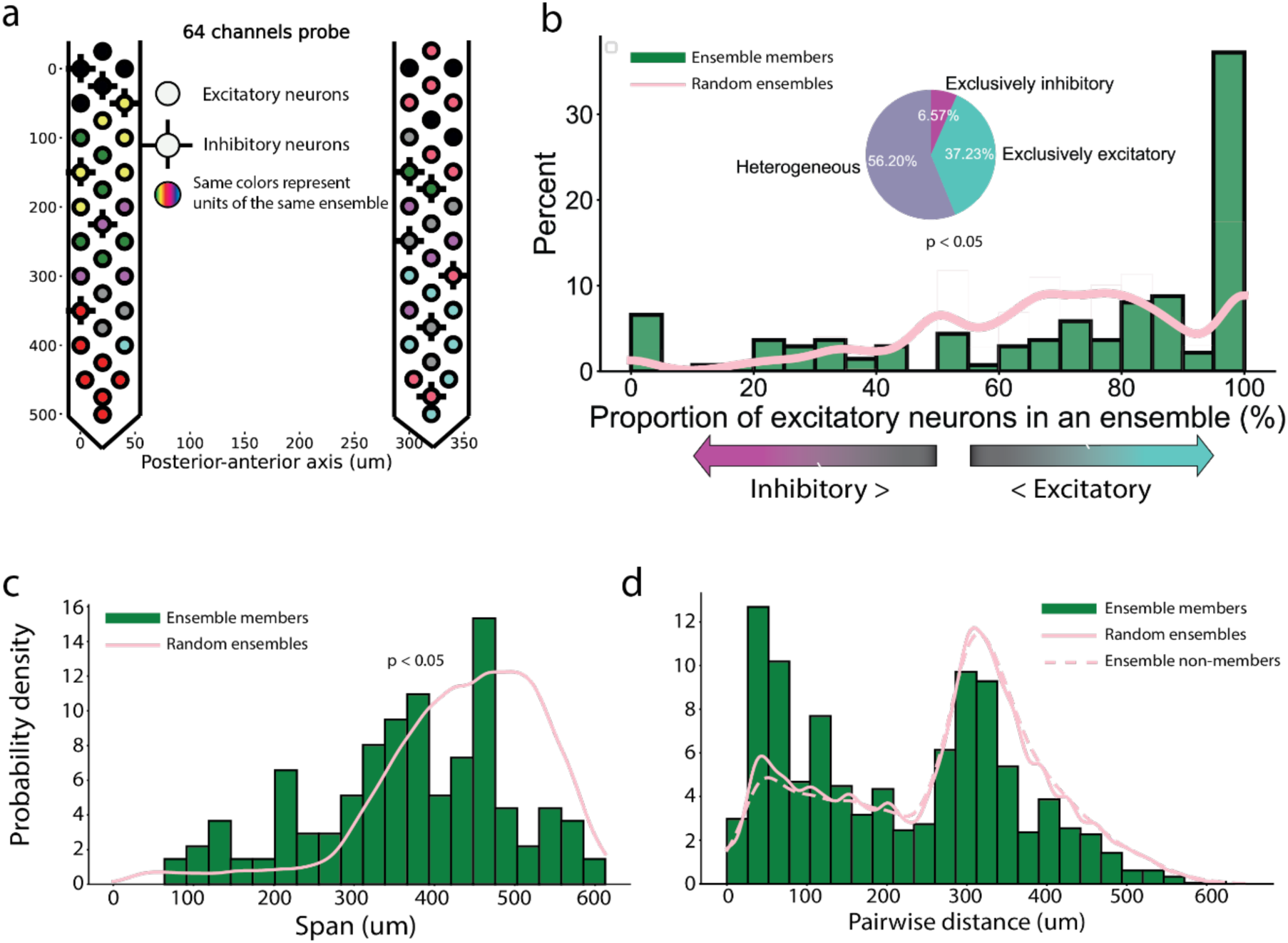
Ensembles composition and distribution. On the histograms, green bars represent real ensemble distribution, the pink lines represent the detection of randomly detected ensembles by simulating 5,000 ensembles that are based on the similar number of members as in the real ensembles. Dashed pink line represents non-members pairwise distance. **a)** Exemplar of ensembles detected in one recording projected into the silicon probes. **b)** Proportion of excitatory cells in ensembles. 100% means only excitatory cells, 0% only inhibitory cells. Most ensembles are heterogeneous (56.20%) **c)** Ensemble span is defined as the maximum distance between two members of the same ensemble. **d)** Pairwise distance between neurons in the same ensemble. The Kolmogorov–Smirnov test was used to compare the different distributions.

### Ensemble adaptation, stability, and selectivity

Do songbird auditory ensembles simply reflect correlated activity patterns? Given that ensembles are defined by the correlated activity of neurons and that most NCM ensembles are biased towards excitatory members, we examined the pairwise correlation between cell types (Figure 5a). Inhibitory neurons typically exhibit higher correlated activity than excitatory neurons in many systems. However, considering that there are fewer inhibitory members in ensembles than expected by chance, one might reason that inhibitory neurons in our data may show weaker pairwise correlations, leading to their under-representation in the detected ensembles (Figure 5a). We found that the overall correlation among inhibitory neurons was higher than that for excitatory units (β= 0.07, se = 0.0006, z = -114.82, p < 0.001, Figure 5b). Also, we confirmed that ensemble members had higher correlation values compared to non-comembers (ý= 0.012, se = 0.0009, z = -13.10, p < 0.001, Figure 5b). Since different cell types have different baseline firing rates, we reasoned that the increased correlation values in inhibitory cells is, on average, due to their higher firing rates. To address this question, we circularly shuffled the data to preserve each neuron’s firing structure, creating a shuffled correlation distribution (Figure 5c). The correlation distribution of inhibitory-inhibitory, inhibitory-excitatory, and excitatory-excitatory pairs of the observed data differed significantly from each other (Figure 5c, bottom), while the circularly-shuffled data have significant overlap in the correlation distribution (Figure 5c, top). Therefore, we observed that even though inhibitory neurons have generally high pairwise correlation values, this is not explained by higher firing rates alone and does not predispose NCM ensembles toward more inhibitory cell membership. More parsimonious is that NCM ensembles share coincident inputs and intra-ensemble synaptic connections among members, regardless of their heterogeneous cellular composition.

**Figure 5.**
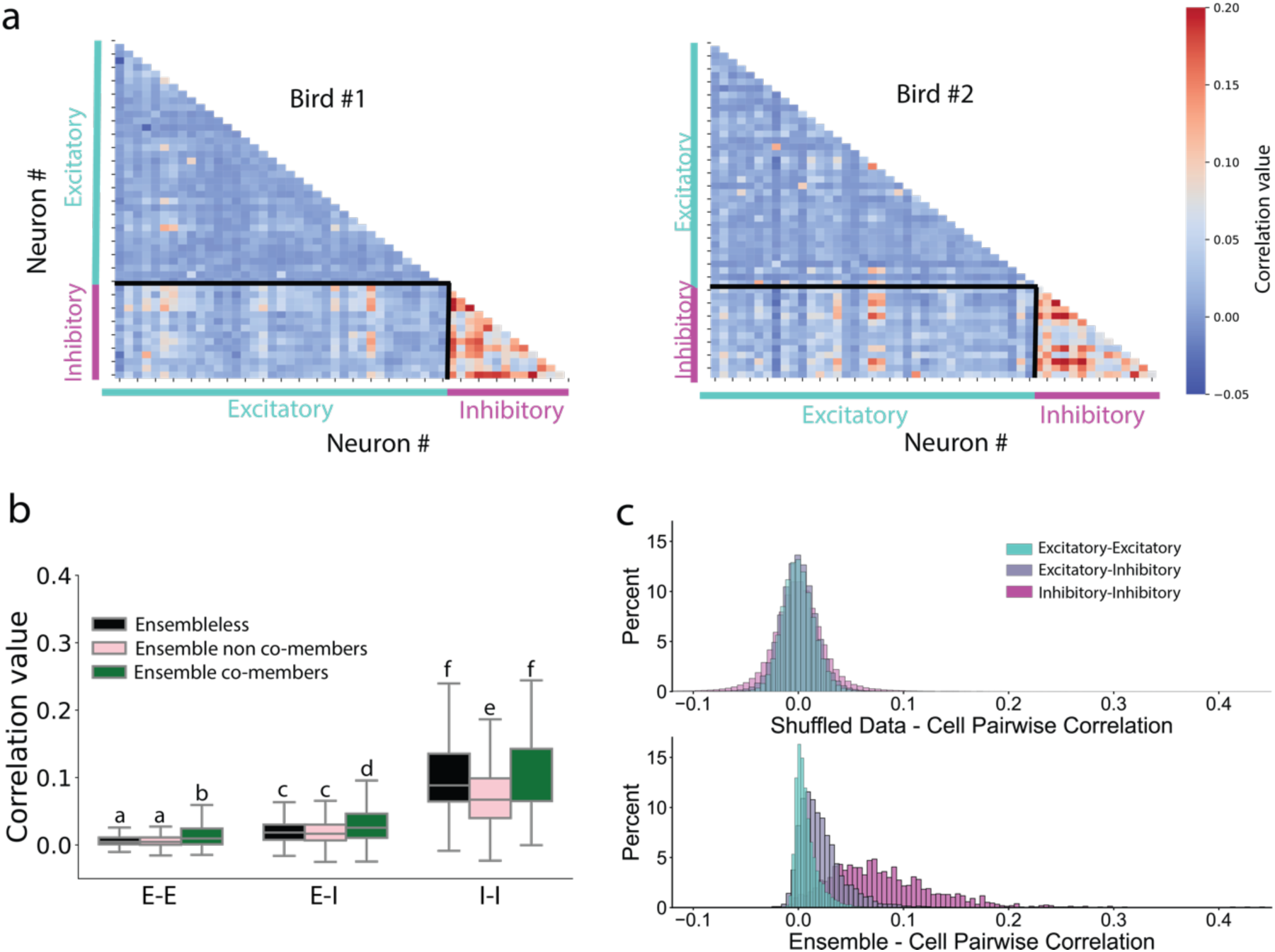
Pairwise spike correlations are higher among inhibitory neurons and ensemble members. **a)** Exemplar of binned spikes pairwise correlation matrix between neurons from two birds. **b)** Correlation between excitatory–excitatory (E-E), excitatory–inhibitory (E-I), and inhibitory–inhibitory (I-I) neuron pairs for ensemble co-members, ensemble non-co-members, and ensembleless members. Letters above boxplots show contrasts within treatment groups with p<0.05 in Tukey’s post hoc test. **c)** Top panel: distribution of 1,500 shuffled data correlation. Bottom panel: observed data pairwise correlation.

### Stimulus-specific adaptation and ensemble dynamics

One key feature of higher-order sensory pallial structures like the songbird NCM is that individual neurons display SSA when presented with successive stimulus repeats (Figure 6a). This indicates that the NCM tracks recently heard songs and responds robustly to novel songs, a feature considered to reflect learning and novel stimulus recognition. We found that excitatory units have a steeper adaptation profile compared to inhibitory units, as measured by the regression slope for response by trial (ks = 0.13, p-value = 0.034; Figure 6b). Our results show that ensembles also exhibit this adaptation feature (Figure 6b) and have adaptation that is similar to inhibitory units and excitatory units (Ensemble – Inhibitory: ks = 0.13, p-value = 0.20; Ensemble – Excitatory: ks = 0.11, p-value = 0.17). Therefore, ensembles, like their constituent members, exhibit robust online recognition of recently-heard sensory experience.

**Figure 6.**
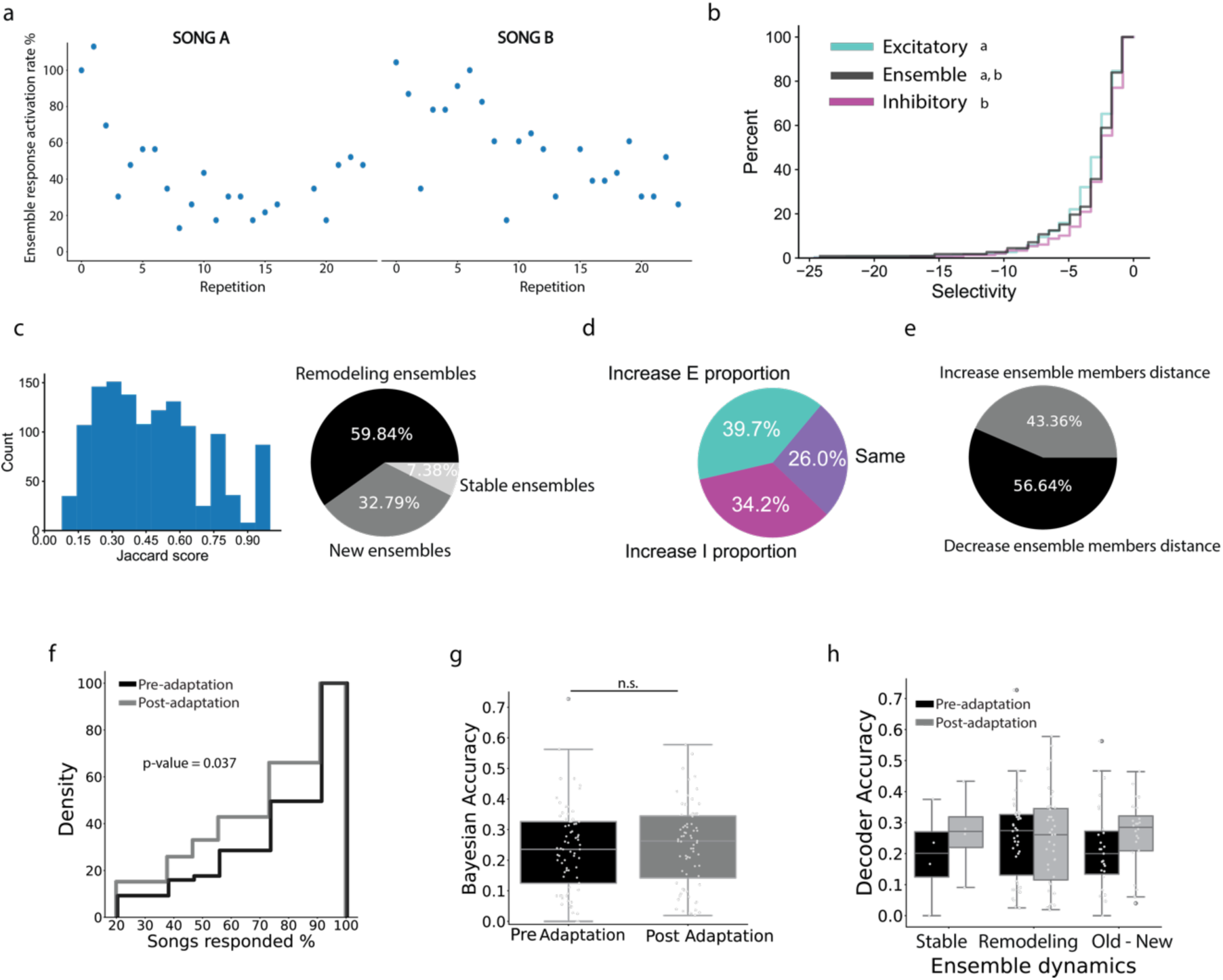
Ensemble dynamics pre- and post-adaptation. **a)** Example of SSA slopes using a linear regression for one cell to two different conspecific songs. This is from unpublished data that played the songs in order and not randomly to identify SSA. **b)** Cumulative distribution function comparing excitatory, inhibitory and ensemble selectivity and their adaptation slope only for adapting neurons, the more negative the slope, the steeper the adaptation. Letters indicate statistical difference between groups. **c)** Jaccard score distribution and the proportion of ensemble dynamics pre- and post-adaptation based on the Jaccard Score. **d)** Change in ensemble composition for new and remodeling ensembles together. **e)** Proportion of ensembles that increased or decreased the pairwise distance between ensemble members pre- and post-adaptation. **f)** Ensemble selectivity measured by the number of songs that the ensembles responded to. The Komologrov-Smirnov test was used to measure the statistical difference. **g)** Comparison of pre- vs post-SSA ensembles decoding accuracy. **h)** Comparison of pre- vs post-SSA ensembles decoding accuracy by ensemble dynamics.

How stable is ensemble membership composition over the course of stimulus adaptation? Adaptation occurred over a relatively short period, approximately 25 minutes, during random playback of 8 different auditory stimuli (Figure 6a, b). We aimed to determine whether ensembles remain stable or undergo changes during this period of adaptation. We identified three patterns of ensemble activation in the recordings (Supplementary figure 3a): 1) stable ensembles, which did not change their neuronal composition throughout the recording; 2) strengthening ensembles, where ensemble members showed increased co-firing; and 3) remodeling ensembles, where one or more members of the ensemble stopped firing in synchrony with the rest. Despite the rapid adaptation, most ensembles remained stable over time, maintaining a consistent composition, with only a few undergoing rapid remodeling or increased connectivity (Supplementary figure 3b). The proportion of inhibitory/excitatory neurons did not differ across these categories of activation pattern (Supplementary figure 3c). Therefore, most ensembles keep track of recently-heard stimuli using SSA in a similar trajectory as their underlying single units, and ensemble neuronal composition is stable through the SSA period. The above analysis could inherently favor detecting membership stability, because the ensemble detection relies on correlated activity during the entire recording. To address this potential bias, we divided the recording into two halves — before (the first 13 trials) and after adaptation (the last 12 trials) — and performed ensemble detection on each segment. After identifying ensembles, we compared those detected in the first and second halves, and ensemble membership was then compared using the Jaccard Similarity Index: If the index was higher than 0.9, the ensembles were defined as stable (7.38% of the ensembles – Figure 6a). If the index was greater than or equal to 0.33 and smaller than 0.9, the ensembles were considered the same but remodeled (59.84% of the ensembles – Figure 6b). If the index was below 0.33, the ensemble was considered entirely new, with the original ensemble no longer existing (32.79% of the ensembles – Figure 6b). Therefore, NCM ensembles showed marked plasticity in their membership over the course of these recordings.

Next, considering the fast remodeling of ensembles during SSA, we investigated whether these changes were driven by specific cell types. We found that post SSA, 38.05% of the ensembles showed an increase in the proportion of inhibitory cells, 32.74% of the ensembles increase in excitatory members, and 29.20% didn’t change the proportion of excitatory/inhibitory cells. Furthermore, we examined whether the members of remodeled ensembles were physically closer or farther apart than the members of ensembles before adaptation. We found that most ensemble members, after adaptation, were detected in channels that were more proximal than before adaptation (56.64% - Figure 6d). In summary, when separating the recording into two halves, we find that only a small number of ensembles stay stable through adaptation. Most ensembles, after adaptation, either have remodeled or new ensembles have formed. The newly formed ensembles showed an increase in the participation of inhibitory neurons, and membership favors shorter physical distances between neurons.

Lastly, given that SSA may reflect a form of cellular memory, we examined the ability of neuronal ensembles to discriminate between distinct auditory stimuli, before and after adaptation. Post-adaptation, the song selectivity for ensembles was significantly enhanced (p = 0.03 – Figure 6f), consistent with increased tuning specificity following familiarity with the recent stimulus environment. However, decoding accuracy (i.e. the ability of ensembles to distinguish one stimulus from another), assessed via a Bayesian classifier, did not differ significantly from pre- to post-SSA (Figure 6g). This lack of change in decoding performance held across remodeling, stable, and newly recruited ensembles (Figure 6h). While both stable and newly formed ensembles exhibited a trend toward improved classification accuracy, these differences did not reach statistical significance. Together, these observations indicate that, despite remodeling, adaptation, and dynamics in selectivity, the ability of ensembles to carry stable stimulus-related activity is well preserved over time.

## Discussion

Neuronal ensembles have been shown to play a crucial role in sensory encoding and memory formation, in particular via increasing the extraction of stimulus signal-to-noise ratio compared to the activity of individual neurons ^10,62,63^. However, some of their fundamental properties remain unclear. In this study, we provide insights into several of these properties. Coordinated neuronal ensembles can be stable over different time scales, ranging from a few minutes up to several weeks ^64^. Nevertheless, if exposed to a new environment or learning regime, the ensemble representation can drift and change composition ^64^. Our results show that most ensembles in a songbird pallial region undergo fast remodeling during the short acquisition phase of SSA and that they become more selective post adaptation. It is likely, therefore, that ensembles recruit co-activated neurons to efficiently encode stimuli and integrate newly learned sounds with previously existing networks that represent familiar sounds.

Although ensembles of inhibitory neurons have been suggested to coordinate the activity of excitatory neurons ^65–70^, there is a gap in the understanding of how both excitatory and inhibitory neurons contribute to ensemble membership and co-membership. Our findings indicate that most NCM ensembles are biased toward excitatory members, more than expected by chance. In NCM, the latency to first spike after stimulus onset is shorter for inhibitory neurons than for excitatory neurons ^27,71^. This feed-forward network architecture, combined with the predominance of excitatory members in ensembles, supports the idea that a small population of inhibitory neurons can coordinate excitatory activity within ensembles.

Another key aspect in understanding the nature and function of neuronal ensembles is the timing of their coordinated activity ^62^. There is significant variability in the temporal frames used to study ensembles, stemming from methodological constraints (e.g., electrophysiology vs. calcium imaging), analytical approaches (e.g., functional ensembles defined by activity during stimulus playback or spike timing within one LFP oscillation), and brain region / species. Most ensemble timing studies have been performed in rodents and employ time windows between 10 ms and 50 ms to analyze the coordinated activity among neurons ^72,73,73–75^. While ensembles comprised exclusively of PV neurons have been shown to synchronize at 2.4 ms ^76^, here, we show that ensembles composed of both inhibitory and excitatory neurons in songbirds are coordinated within a very fast time window of < 10 ms. This rapid neuronal synchronization supports the idea that songbirds perceive their auditory world in a finer temporal frame than many mammalian species ^77^, and is likely key to their rapid vocal communication signaling.

The rapid coordination of NCM ensembles (6ms), combined with the observation that most neurons are part of only one ensemble, suggests that in the songbird pallium contains fast synaptic connections that coordinate this activity. This coordination could be due to chemical or electrical synapses, given that gap junction proteins are widely expressed in the songbird pallium ^78^. Understanding the fast ensemble coordination via electrical synapses could be important to unpacking highly time-variant stimuli such as song with high temporal precision.

In conclusion, we provide evidence that during stimulus-specific adaptation, the local neuronal network undergoes restructuring. Additionally, we demonstrate that excitatory and inhibitory neurons can be part of the same neuronal ensemble and that the time frame for ensemble synchronization in songbirds is exceptionally fast.

### Limitation and future directions

Although we classified the cells as inhibitory or excitatory, we lack detailed information about their subtypes, such parvalbumin or somatostatin interneurons ^79,80^. Thus, it remains unclear which inhibitory or excitatory cell subtypes contribute to their composition. Regarding the timing of ensembles, while we identified 6ms as potentially the optimal bin size, different ensembles may be synchronized on different time scales. Some ensembles might exhibit very fast coordinated activity, while others—particularly those composed exclusively of excitatory cells—might synchronize more slowly. Future experiments could systematically examine different optimum time bins to help resolve this question. The ensemble dynamics we report here are consistent with short term stability or remodeling and future studies could investigate how social communication context and neuromodulators shape NCM ensemble dynamics. Although the unsupervised model that combines PCA with ICA that we used here is widely used ^41,45,72,81,82^, the threshold ensemble activity and ensemble membership is determined by null hypothesis distribution which has not been linked to physiological phenomena ^10^. Other methods have been used to identify temporally clustered neuronal activity as ensembles, such as coactivity that happens within sharp wave ripple oscillations ^76^, pattern completion based activation ^83^, and time sequence activity patterns ^6^.

## Acknowledgments

We thank members of the Healey lab for helpful comments on previous versions of this manuscript. We also thank the Animal Care Staff and many undergraduate assistants at the University of Massachusetts Amherst for taking care of our feathered friends. Support for this work from NIH R01NS082179 (LRH).

---

**Supplementary figure 1.**
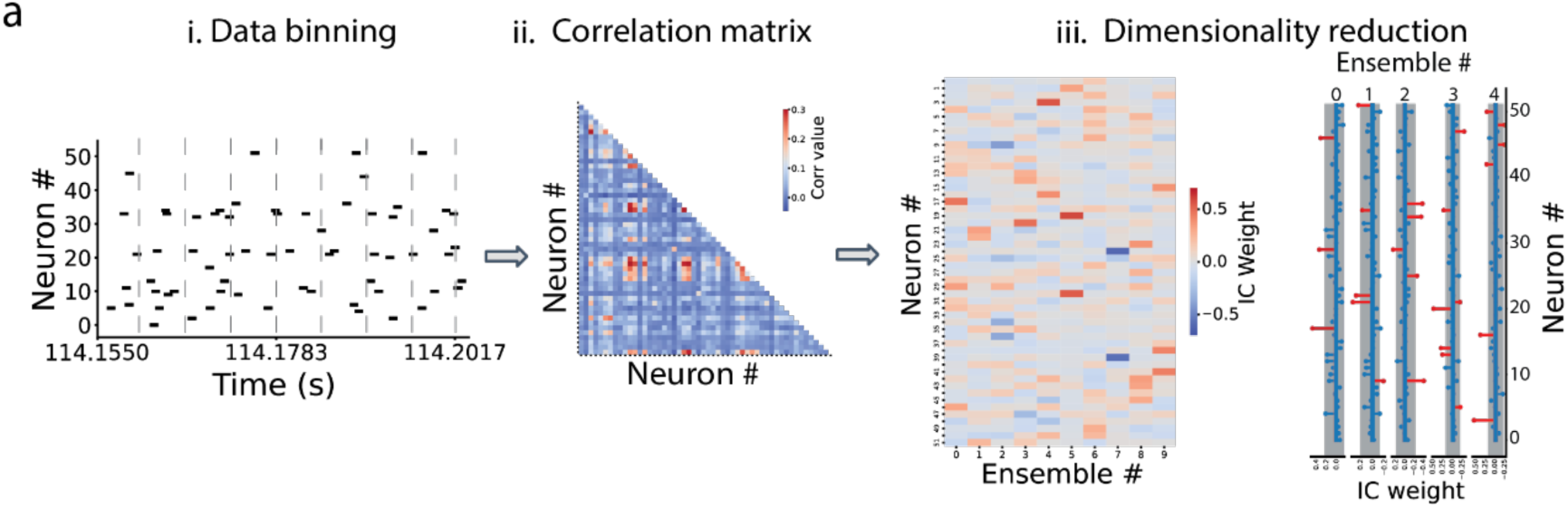
**a)** Procedure for ensemble detection. *i)* Data binning of z-scored data (different bins tested – from 2 – 100 ms). ii) Correlation matrix of binned spike trains. Independent components (IC) weights of neurons for each detected ensemble. *iii)* Ensemble members (red stems) have IC weights exceeding the threshold shown as grey areas.

**Supplementary figure 2.**
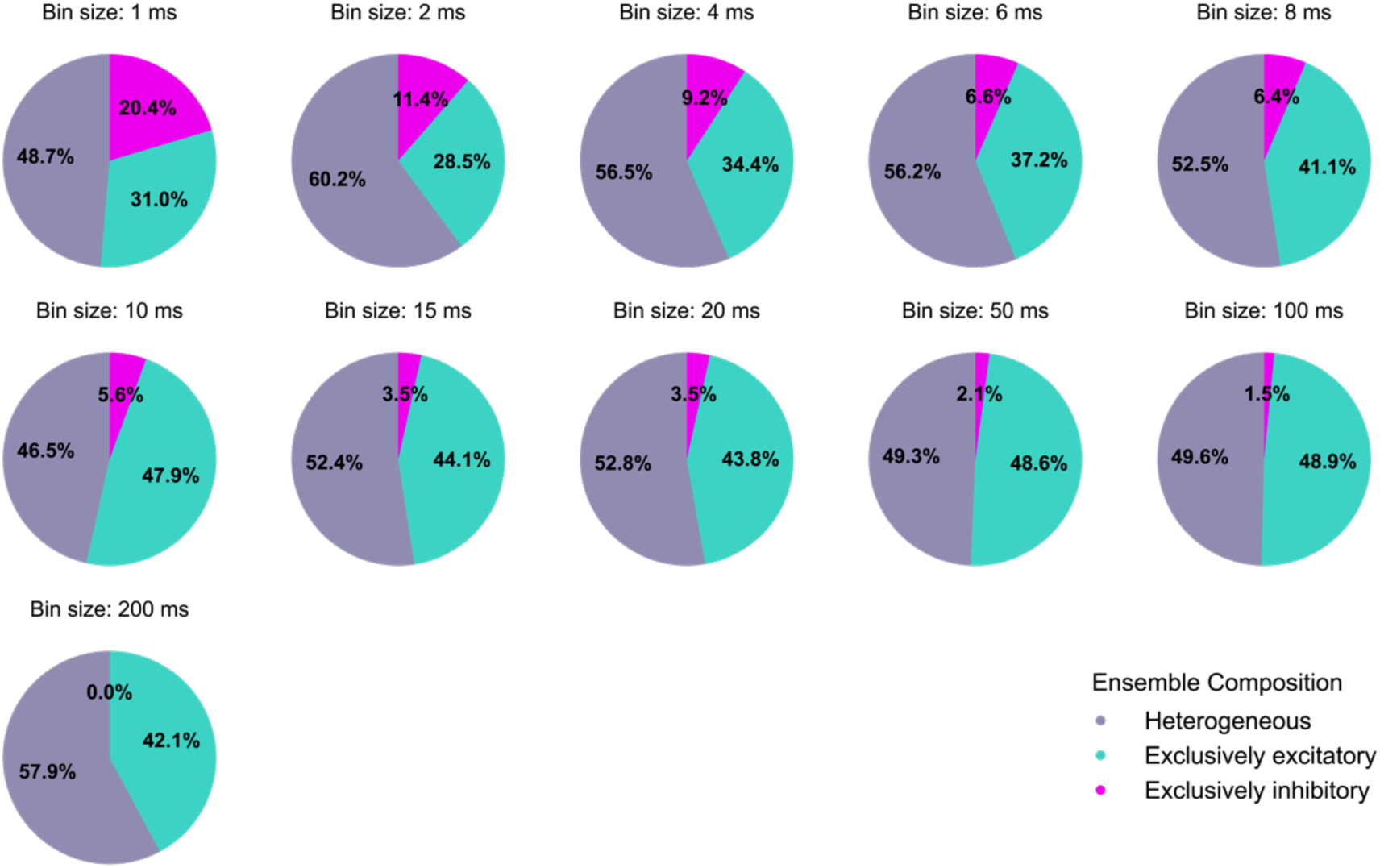
Ensemble composition by bin size. As bin sizes get larger there is a decrease in the proportion of exclusive inhibitory ensembles.

**Supplementary figure 3.**
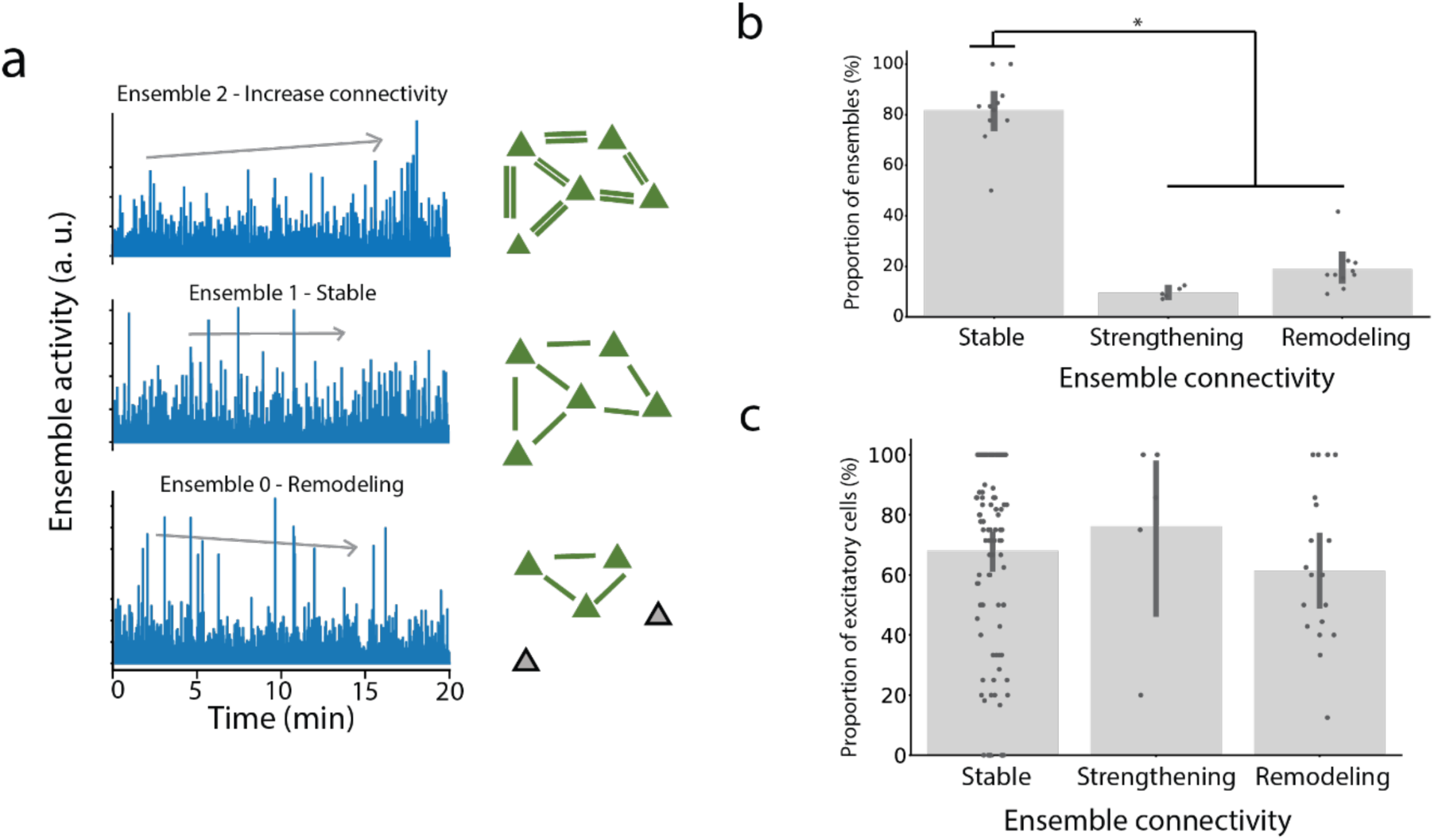
**a)** *Top left*: One example “strengthening” ensemble. *Middle left:* One example “*stable*” ensemble. *Bottom left,* One example “*remodeling*” ensemble. ***p<0.001 Mann-Kendall trend test. On the *right,* schematics representations of what each ensemble dynamics look like. **b)** Proportion of ensembles classified in the three possible different connectivity dynamics. **c)** Proportion of excitatory cell in each ensemble dynamics. Tukey’s post hoc test results are shown at the bottom. * p <0.05, n.s. no significant difference

